# A non-lytic membrane permeabilization program drives epithelial cell turnover *in vivo*

**DOI:** 10.64898/2026.07.06.736761

**Authors:** Motohiro Morikawa, Sa Kan Yoo

## Abstract

A central dilemma of epithelial cell turnover is eliminating and replacing cells while simultaneously preserving tissue architecture and barrier function. Conventionally, apoptotic or non-apoptotic cell extrusion has been implicated in the intestinal epithelial turnover. Here, we identify a non-lytic membrane permeabilization program that drives physiological enterocyte turnover *in vivo*. In the *Drosophila* intestine, enterocytes undergo erebosis, a non-apoptotic form of cell death characterized by depletion of cytoplasmic proteins. We discover that this process is mediated by transient plasma membrane pores with estimated diameters of 16–50 nm, permitting extracellular protein influx and loss of cytoplasmic contents. The pore-forming protein Ninjurin A (NijA) accumulates as puncta during erebosis, and is necessary and sufficient for driving this process. NijA-mediated transient permeabilization preserves the membrane framework of dying cells, enabling their replacement without disrupting epithelial barrier architecture.

---

Under homeostatic conditions, cells in many organs exist in a constant state of flux. The intestinal epithelium has the fastest cell turnover rate (Sender and Milo, 2021). Traditionally, apoptotic or non-apoptotic cell extrusion has been implicated in the intestinal epithelial turnover (Bullen et al., 2006; Ghazavi et al., 2022; Krueger et al., 2025). Recently, we demonstrated that, in the *Drosophila* gut, homeostatic cell turnover under physiological conditions is not regulated by apoptosis or lytic cell death such as necrosis (Ciesielski et al., 2022). Apical extrusion does not mediate it either. Rather, an enigmatic phenomenon, termed erebosis, is the dominant mechanism of fly gut cell turnover (Ciesielski *et al*., 2022). Erebosis is a process in which cells lose essential components and accumulate intracellular Ance in *Drosophila* enterocytes. Erebosis is distinct from apoptosis, necrosis and autophagy. The mechanisms underlying this phenomenon, whether involving vesicle trafficking, protein degradation, secretion or excretion, reduced transcription or translation, or membrane pore formation, have remained unclear.

Previously, we demonstrated that the loss of intracellular GFP and accumulation of Ance occur with closely correlated timing (Ciesielski *et al*., 2022). In addition, a small amount of Ance is normally present above enterocytes in the gut lumen, the spatial patterns of Ance mRNA expression and protein localization differ, and secretion of Ance is necessary for its intracellular accumulation in erebotic cells (Ciesielski *et al*., 2022). Together, these observations led us to hypothesize parsimoniously that transient membrane pores form at the initiation of erebosis, allowing diffusive exchange of extracellular and intracellular molecules. On the contrary to a variety of necrotic cell death, such as necrosis, necroptosis and pyroptosis, involving membrane pore formation and cell lysis (Galluzzi et al., 2018; Kayagaki et al., 2015; Yuan and Ofengeim, 2024), erebotic cells are not necrotic at the very time of imaging: addition of propidium iodide (PI) (Ciesielski *et al*., 2022) or Sytox does not label erebotic cells (Fig 1A), indicating that erebotic cells retain intact plasma membrane integrity. Thus, if formed, membrane pores should be closed immediately after opening.

**Figure 1.**
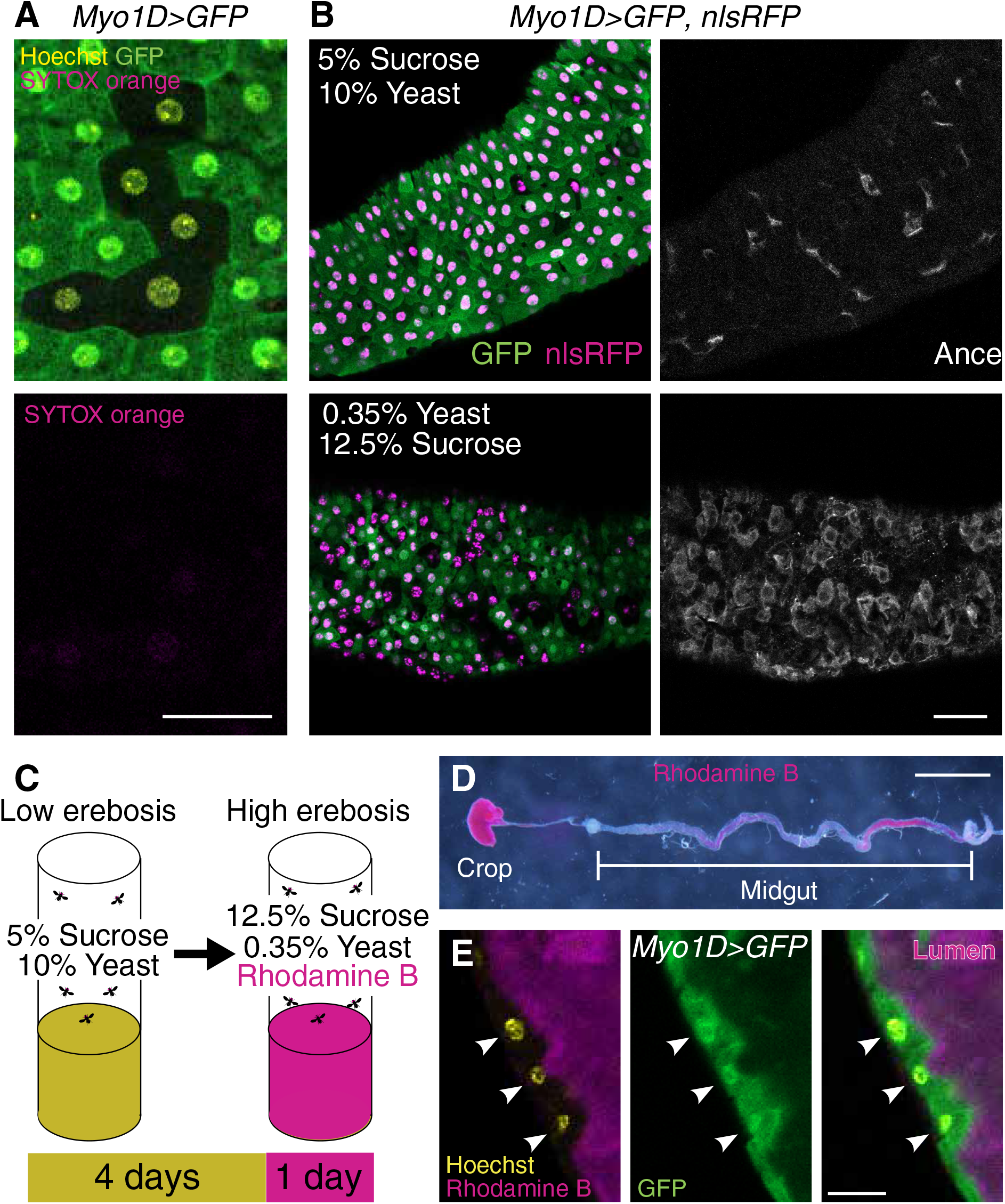
Erebosis induction and Rhodamine feeding. (A) Erebotic cells are not necrotic. GFP (-) erebotic cells are not stained by membrane-impermeable SYTOX orange (magenta). Scale bar, 40 μm. (B) Low-yeast, high-sugar food induces erebosis. High-yeast, low-sugar 1x SY food (top) results in only a few GFP (-) nlsRFP (+) erebotic cells, while low yeast high sugar food (bottom) leads to many erebotic cells. In both conditions, GFP (-) erebotic cells accumulate Ance. Scale bar, 50 μm. (C) The scheme of acute erebosis induction. Erebosis was suppressed for 4 days followed by acute induction by low-yeast, high-sugar food. 0.1% Rhodamine B was added to the low-yeast, high-sugar food to label erebotic cells. (D) Ingested Rhodamine (red) localizes in the lumen of the crop and the midgut. Scale bar, 1 mm. (E) Rhodamine (magenta) localizes to the gut lumen, while it is not incorporated into normal ECs expressing GFP (arrowheads). Scale bar, 20 μm.

If pores are truly formed, we reasoned that cell-impermeable tracer dyes present during erebosis could label cells, while failing to label cells that have already undergone erebosis at the time of imaging. We investigated whether prior feeding of fluorescent tracer dyes before imaging would label erebotic cells. To efficiently label erebotic cells at the time of their occurrence and to ensure fluorescent dyes are present during erebosis, we first sought a condition that induces erebosis. We detect erebotic cells based on intracellular accumulation of Ance and/or the reduction of fluorescent protein signals in enterocytes of the R4 region (Ciesielski *et al*., 2022). Through screening conditions that promote erebosis, we found that a high-yeast, low-sugar diet suppresses erebosis, while a low-yeast, high-sugar diet enhances erebosis, even though both conditions have the same caloric content (Fig 1B). We developed a scheme where flies were kept in an erebosis-suppressive condition (high yeast/low sugar) for 4 days and then incubated with the fluorescent dye Rhodamine B under the erebosis-inducing condition (low yeast/high sugar) for 1 day (Fig 1C). Fed Rhodamine B was readily observed in the lumen but not within the gut tissue layer (Fig 1D-E).

We found that prefeeding Rhodamine B labels erebotic cells (Fig 2A). The labeling pattern was diffuse in the cytoplasm, strongly negating the possibility of endocytosis mediating this process. Endocytosed dye would be expected to appear in vesicular structures (Allen et al., 2020; Gupta et al., 2009; Sriram et al., 2003) rather than diffusely throughout the cytoplasm. Erebotic cells pre-labelled by Rhodamine B were not labeled by another non-cell permeable tracer, a fluorescent phalloidin, added at the very time of imaging, indicating that they possess the normal impermeable plasma membranes, not being necrotic (Fig 2B). Furthermore, Rhodamine B+, GFP-erebotic cells retained nlsRFP, also indicating that they are distinct from necrosis. Time-course analysis demonstrates that Rhodamine B begins to label erebotic cells 4 hours after feeding (Fig 2C-D). There were also erebotic cells that were not labelled by Rhodamine B even at 4 or 18 hours post feeding, reflecting cells that had been already erebotic when Rhodamine B reached enterocytes. This indicates that erebotic cells can remain in the tissue likely to maintain the barrier function after its initial process of losing cytoplasmic GFP. Consistent with our previous findings, 48 hours post feeding, some Rhodamine B-positive erebotic cells were observed as flattened cells positioned above newly emerging enterocytes (Fig S1), which contrasts with apically extruded round apoptotic cells.

**Figure 2.**
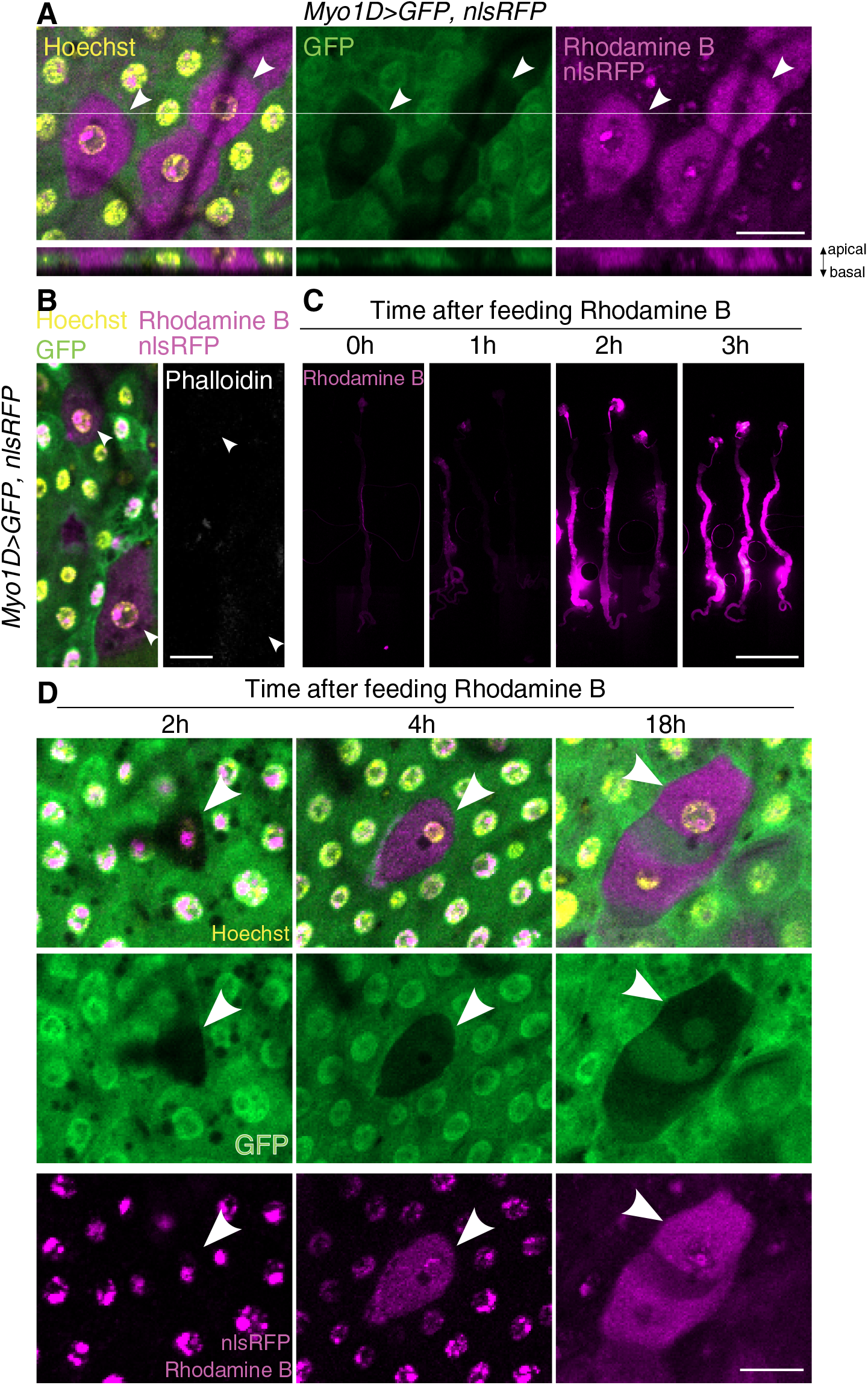
Erebotic cells are labeled by pre-fed rhodamine B. (A) After the erebosis induction and 0.1% Rhodamine B feeding, GFP (-) erebotic cells (arrowheads) become Rhodamine (+). The bottom pictures show Z projection of GFP (+) normal ECs and GFP low erebotic cells. Rhodamine is exclusively incorporated into erebotic cells. Scale bar, 20 μm. (B) GFP (-) or low erebotic cells (arrowheads) are Rhodamine (+) but not stained with membrane impermeable phalloidin (white). Scale bar, 20 μm (C) Rhodamine enters the lumen of all guts 2 hours after Rhodamine feeding. Scale bar, 2 mm. (D) GFP low or (-) erebotic cells (arrowheads) become Rhodamine (+) 4 hours after Rhodamine feeding. Scale bar, 20 μm.

We rigorously tested our hypothesis that the transient membrane pores account for the Ance accumulation and the loss of cytoplasmic GFP. If temporary membrane pores form during erebosis, there should be a size limit for the tracer dye that can enter erebotic cells. We used two different sizes of fluorescent dyes, TRITC (Rhodamine B derivative) conjugated with dextran: 150 kDa dextran-TRITC and 2000 kDa dextran-TRITC. These are larger than 71 kDa Ance and 27 kDa GFP. The 150 kDa dextran-TRITC labeled erebotic cells, but the 2000 kDa dextran-TRITC did not (Fig 3A), although both dyes could reach the apical edge of enterocytes (Fig S2). The data that the 150 kDa tracer can enter cells are consistent with the idea that membrane pores allow the passage of Ance and GFP. Besides the diffuse pattern of the dyes, the existence of the size limit for tracer dyes to enter cells during erebosis strongly supports the temporary formation of membrane pores, but not endocytosis or exocytosis. The hydrodynamic diameter of 150 kDa dextran is approximately 16 nm, while that of 2000 kDa dextran is around 50 nm (Chen et al., 2022). Thus, the pore diameter is likely between 16 and 50 nm.

**Figure 3.**
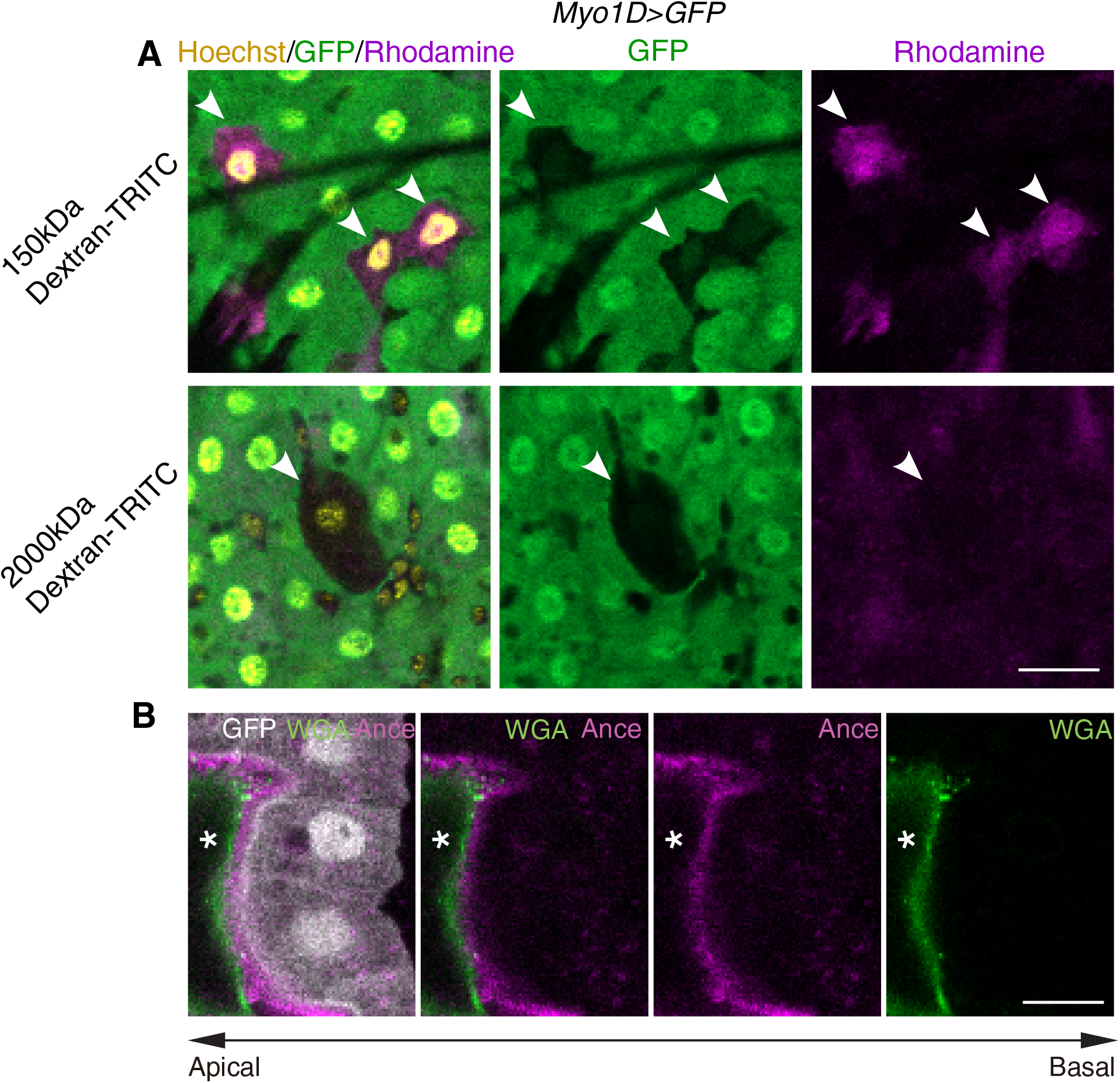
Incorporation of fluorescent tracers is size-dependent. (A) GFP (-) erebotic cells (arrowheads) are 150 kDa dextran-TRITC (+) (top) but 2000 kDa dextran-TRITC (-) (bottom). Scale bar, 20 μm. (B) Ance (magenta) localizes just above enterocytes (GFP, white) and below the peritrophic matrix (WGA, green). White asterisks indicate the gut lumen. Scale bar, 10 μm.

Our data strongly suggest that there is a transient pore formation during erebosis. This can simply explain how cytoplasmic protein such as GFP diffuses out of cells and extracellular Rhodamine B, which are fed and abundantly exists in the lumen, diffuses into cells, but still raises the question of how Ance, which accumulates at the apical surface, can enter enterocytes: Ance needs to exist abundantly above enterocytes but must not strongly bind to enterocytes in order to be able to diffuse into cells. To elucidate how Ance enables this in the lumen, we examined its exact location at the apical surface of enterocytes. We analyzed spatial relationships among enterocytes, Ance and the peritrophic membrane. The peritrophic membrane is a thin layer composed of chitin and glycoproteins that covers the apical side of enterocytes (Hegedus et al., 2009; Kuraishi et al., 2011; Lehane, 1997), which can be labeled by fluorescent wheat germ agglutinin (WGA) (Katheder et al., 2023; Pandey et al., 2023). We found that Ance fills the interspace between enterocytes and the peritrophic membrane (Fig 3B). This accumulation of Ance filling the interspace would allow its influx into enterocytes during erebosis.

What is the molecular mechanism underlying transient pore formation? We interrogated molecules that might function as pore-forming proteins, including Attacin-D (Oi et al., 2025), Torso-like (Mineo et al., 2018) and Ninjurin A (NijA) (Broderick et al., 2012; Kayagaki et al., 2021). Notably, flies lack both Gasdermin D and MLKL, which are responsible for pore formation during pyroptosis (Kayagaki *et al*., 2015; Santa Cruz Garcia et al., 2022) and necroptosis in mammals (Cai et al., 2014; Ros et al., 2017), respectively. Among the molecules examined, only NijA knockdown inhibited erebosis (Fig 4A-C). Ninjurins are a conserved family of transmembrane proteins, originally identified through their upregulation in injured nerves(Araki and Milbrandt, 1996), and more recently shown to regulate lytic cell death in mammals (Kayagaki *et al*., 2021). In Drosophila, ectopic expression of NijA induces non-apoptotic cell death in hemocytes (Broderick *et al*., 2012). Mammalian Ninjurin has been proposed to generate membrane pores either through Ninjurin filament-based structures (Degen et al., 2023) or solubilization of Ninjurin-wrapped membrane discs (the “cookie-cutter” model) (David et al., 2024; Ramos et al., 2024).

**Figure 4.**
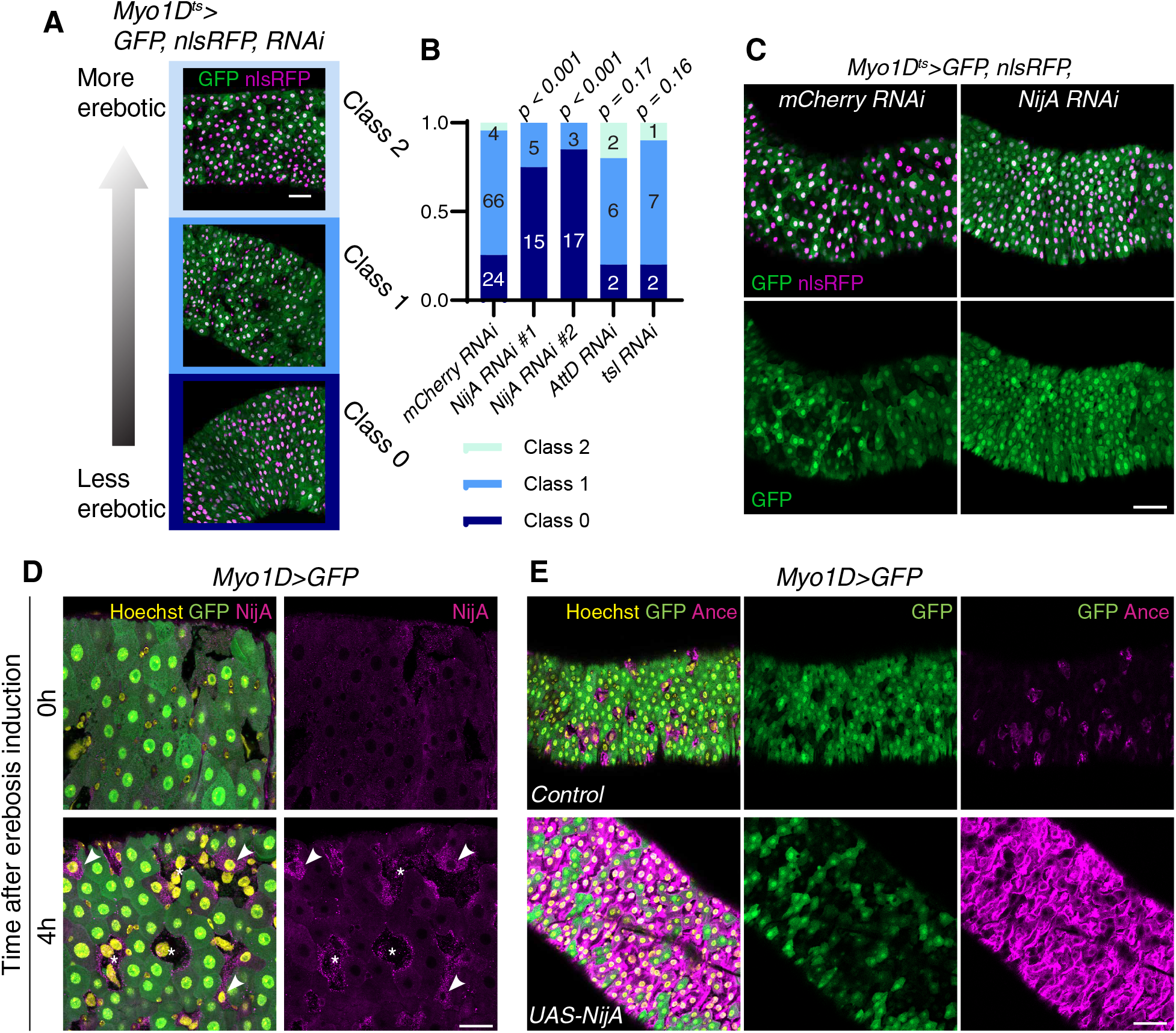
Ninjurin A executes erebosis. (A) Classification method of the frequency of erebosis. GFP and nlsRFP expressions are induced by enterocyte-specific *Myo1D-Gal4* driver. GFP (+) nlsRFP (+) cells are normal enterocytes, while GFP (-) nlsRFP (+) cells are erebotic cells. Class 0 is the least erebotic state with 3 or less erebotic cells/view. Class 1 is mildly erebotic, with less than 30% erebotic cells. Class 2 is massively erebotic, with 30% or more erebotic cells. Scale bar, 50 μm. (B) RNAi screening of pore-forming proteins. Two different *NijA RNAi*s significantly reduce erebosis, while *AttD RNAi* or *tsl RNAi* do not. (C) Representative images of control *mCherry RNAi* and *NijA RNAi* guts. *NijA RNAi* reduces GFP (-) nlsRFP (+) erebotic cells. Scale bar, 50 μm. (D) NijA forms puncta during erebosis. Early erebotic cells with weak GFP (arrowheads) have many NijA puncta while late erebotic cells with much lower GFP signals (asterisks) have fewer puncta. Scale bar, 25 μm. (E) *NijA* overexpression induces erebosis. Compared with control (upper panel), *UAS-NijA* (lower panel) results in massive erebosis with many GFP (-) Ance (+) erebotic cells. Scale bar, 50 μm.

We detected endogenous NijA using an antibody raised against NijA (Broderick *et al*., 2012). Upon erebosis induction, NijA puncta were detected in cells undergoing erebosis with weak GFP signals (Fig 4D). Interestingly, cells at a later stage of erebosis, in which GFP had been completely lost, reduced the number of NijA puncta, suggesting that NijA puncta themselves were removed, consistent with the cookie-cutter model. Finally, we examined whether NijA expression is sufficient to induce erebosis. Ectopic expression of NijA in enterocytes for one day induced massive erebosis, as detected by loss of GFP and accumulation of Ance (Fig 4E).

Here we demonstrate that NijA-mediated transient pore formation is responsible for erebosis (Fig S3). Our data with fluorescent tracer dyes suggest that these transient pores allow the passage of molecules as large as 150 kDa, consistent with entrance of 71 kDa Ance. The estimated pore diameter is between 16 and 50 nm. This estimated size is comparable to the known size of mammalian Ninjurin 1-mediated pore with median of ~35 nm (David *et al*., 2024). The Ninjurin-mediated pore size is slightly larger than ~21 nm pores formed by Gasdermin D during pyroptosis (Santa Cruz Garcia *et al*., 2022) and the ~16 nm pores by Perforin during apoptosis (Law et al., 2010), and much larger than the ~4 nm pores by MLKL oligomers during necroptosis (Ros *et al*., 2017).

Our findings provide a physiological context for transient, sublethal membrane pore formation in vivo. Previous studies have largely examined sublethal membrane permeabilization under artificial conditions, such as detergent treatment, laser-induced injury or mechanical damage (Jimenez et al., 2015; Suda et al., 2024), leaving its physiological relevance under homeostatic conditions unclear. On the other hand, Ninjurin has primarily been implicated in catastrophic plasma membrane rupture under artificially-inflicted stresses (Kayagaki *et al*., 2021; Ramos *et al*., 2024; Zhu et al., 2025). Our findings uncover an unexpected role for Ninjurin in mediating transient, non-lytic membrane permeabilization under physiological conditions in vivo.

A remaining question is how this Ninjurin-mediated pore can be closed in enterocytes, not leading to necrosis. ESCRT-mediated membrane repair might be responsible, since it is known to limit several types of cell death, including necroptosis (Gong et al., 2017), pyroptosis (Ruhl et al., 2018), and ferroptosis (Dai et al., 2020; Pedrera et al., 2021). Notably, *Drosophila* enterocytes can regrow after formation of transient membrane pores artificially generated by pore-forming toxins secreted by Serratia marcescens (Lee et al., 2016; Socha et al., 2023). In the mammalian gut, Gasdermin D-mediated pores can be repaired by ceramide-mediated membrane remodeling (Nozaki et al., 2022). Taken together with our results, these findings suggest that gut enterocytes might be particularly equipped to tolerate and possibly repair transient pore formation. We propose that this transient pore formation allows gut enterocytes to quietly die while maintaining their membrane architecture intact and preserving the barrier function.

## Methods

### Fly husbandry

Flies were maintained as previously described (Sasaki, Nishimura et al. 2021). They were kept on standard fly food containing 0.8% agar, 10% glucose, 4.5% corn flour, 3.72% dry yeast, 0.4% propionic acid, and 0.3% butyl p-hydroxybenzonate. Crosses were maintained at either 18°C or 25°C, and the resulting progeny were shifted to 30°C to induce GFP/nlsRFP expression simultaneously with the transition to 5% sucrose/10% yeast food.

### *Drosophila* stocks

The following stocks were used in this study.

*Oregon R*

*w-; Myo1D-Gal4, UAS-GFP; tub-gal80*^*ts*^ *(MG*^*ts*^)

*w-; Myo1D-Gal4, UAS-GFP, UAS-nlsRFP; tub-gal80*^*ts*^ *(MGR*^*ts*^)

*mCherry RNAi* (BDSC 35785)

*NijA RNAi* (NIG HMC02999)

*NijA RNAi* (VDRC 5208)

*AttD RNAi* (BDSC 55979)

*tsl RNAi* (BDSC 56967)

*UAS-NijA* (A gift from Dr. Andrea Page-McCaw)

### Food conditions

Virgin female flies were collected and put in yeast-rich 1x SY food containing 0.8% agar, 5% sucrose, 10% dry yeast, 0.4% propionic acid, 0.3% butyl p-hydroxybenzonate (Bass 2007 J Gerontol A Biol Sci Med Sci.) or high sugar low yeast food containing 0.8% agar, 12.5% sucrose, 0.35% dry yeast, 0.4% propionic acid, 0.3% butyl p-hydroxybenzonate.

### Dissection, fixation, Ance and PM staining, and observation

Following the feeding of each food for 7 days, guts were dissected in 1xPBS, fixed for 60 minutes at RT in PBS with 4% paraformaldehyde, and washed by PBSTx (1x PBS with 0.1% Triton X-100) 3 times, followed by an incubation in PBSTx 10% Normal goat serum (NGS), 1/1000 rabbit anti-Ance antibody or 1/100 guinea pig anti-NijA antibody at 4 C overnight. After the incubation, guts were washed with PBSTx 3 times and incubated with PBSTx 10% NGS 100 ug/mL Hoechst 33342 (Thermo Fisher Scientific H1399), 1/500 Goat anti-rabbit Alexa Fluor 633 or Goat anti-guinea pig Alexa Fluor 568 (Thermo Fisher Scientific A21070, A11075) with or without 2 ug/mL WGA Alexa Fluor 555 (Thermo Fisher Scientific W32464) at 4 C overnight. Following the incubation, guts were washed with PBSTx 5 ttimes, mounted, and observed with a confocal microscope (LSM900, Zeiss).

### Acute erebosis induction and fluorescent dye feeding

After virgin collection, female flies were fed with 1x SY food for 4 days. Then, to acutely induce erebosis, flies were moved to high sugar low yeast food containing fluorescent dyes (0.1% Rhodamine B (TCI)), 1% 150 kDa Dextran-TRITC or 5% 2000 kDa Dextran-TRITC (TdB Labs)) and kept for 1 day, unless otherwise specified. Concentrations of 150 kDa and 2000 kDa Dextran-TRITC were determined so that they include the same amount of fluorescent TRITC molecules following the manufacturer’s datasheets.

### Dissection, staining, and live imaging

After the fluorescent dye feeding, guts were dissected in PBS and stained with PBS 100 μg/mL Hoechst 33342 with or without 1/100 Alexa Fluor 647 Phalloidin (Thermo Fisher Scientific A22287) at RT for 10 minutes. Following the staining, guts were washed with PBS and mounted in PBS.

### Statistics

Statistical analysis was conducted with R version 4.5.3. P-values were calculated using Fisher’s exact test. Multiple comparisons were adjusted by Benjamini-Hochberg method.

## Acknowledgement

We thank Elwyn Isaac for the Ance antibody, Andrea Page-McCaw for NijA antibody and UAS-NijA stock, and TRiP at Harvard Medical School, the Bloomington Stock Center, National Institute for Genetics, and Vienna Drosophila Stock Center for fly stocks. We thank the Yoo lab members for helpful comments on the manuscript. We thank Noboru Mizushima for suggesting us feeding a fluorescent dye. This work was supported by RIKEN Junior Research Associate Program to M.M. and JST FOREST (JPMJFR216F) to S.K.Y.

## Author Contributions

M.M. and S.K.Y. conceived the project. S.K.Y. oversaw the project. M.M. executed the experiments. M.M. and S.K.Y. wrote the paper.

---

**Figure S1.**
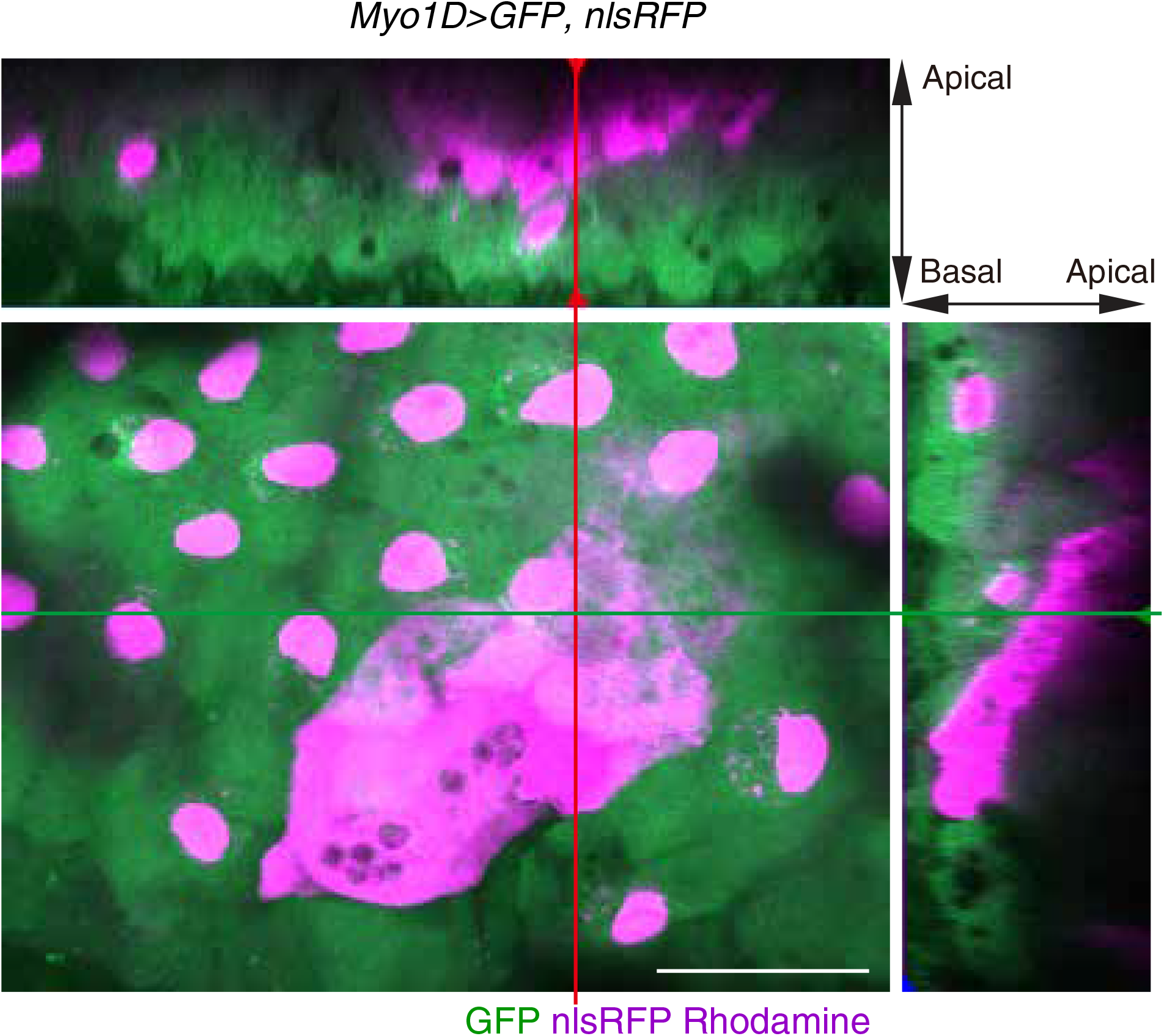
Several erebotic cells localize above newly emerging enterocytes. Rhodamine (+) erebotic cells become flat above the newly emerging enterocytes. Scale bar, 20 μm.

**Figure S2.**
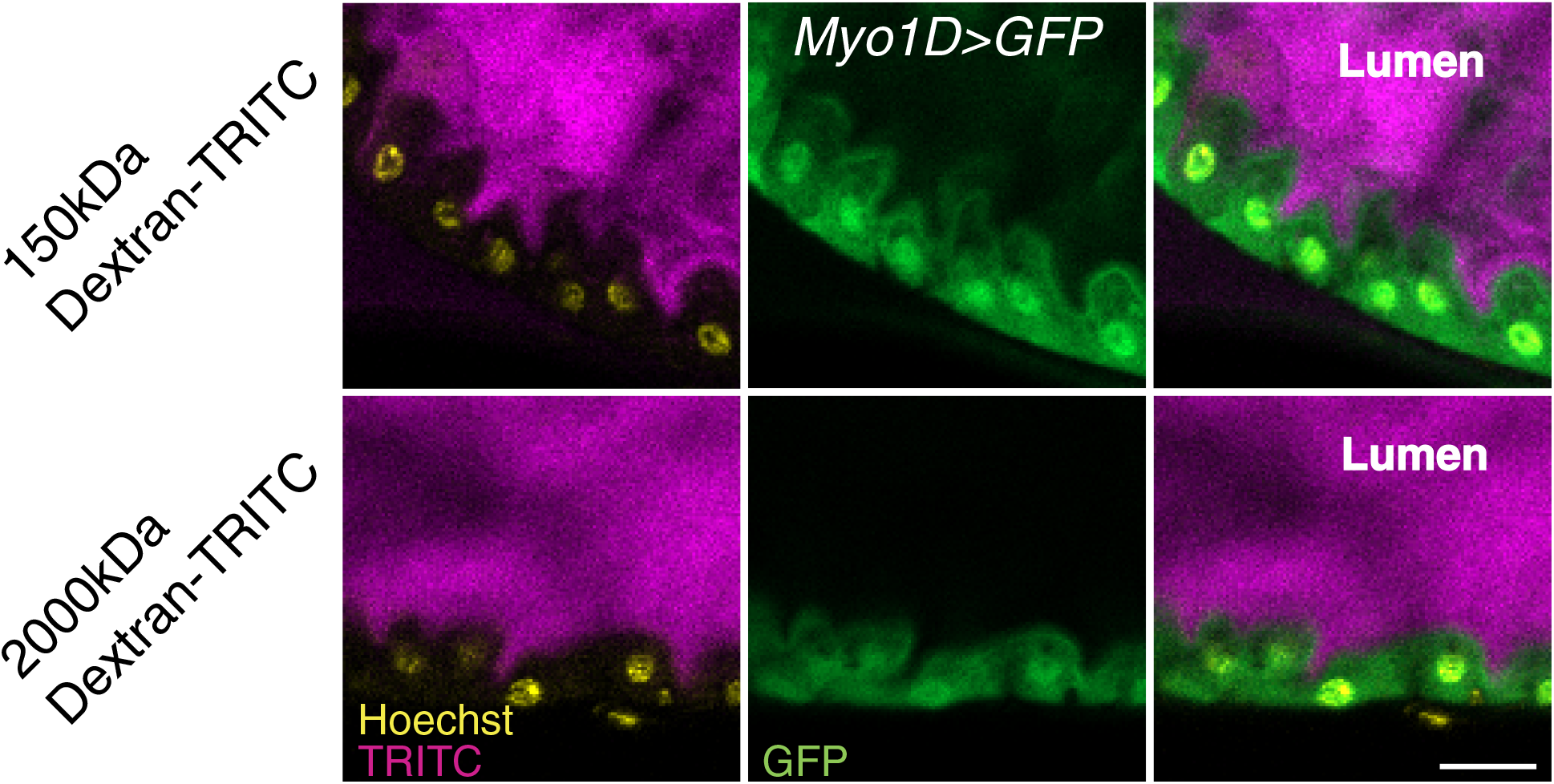
Dextran enters the luminal surface of enterocytes. Both of 150 kDa Dextran-TRITC (upper panel) and 2000 kDa Dextran-TRITC (lower panel) localize to the luminal surface of enterocytes. Scale Bar, 20 μm.

**Figure S3.**
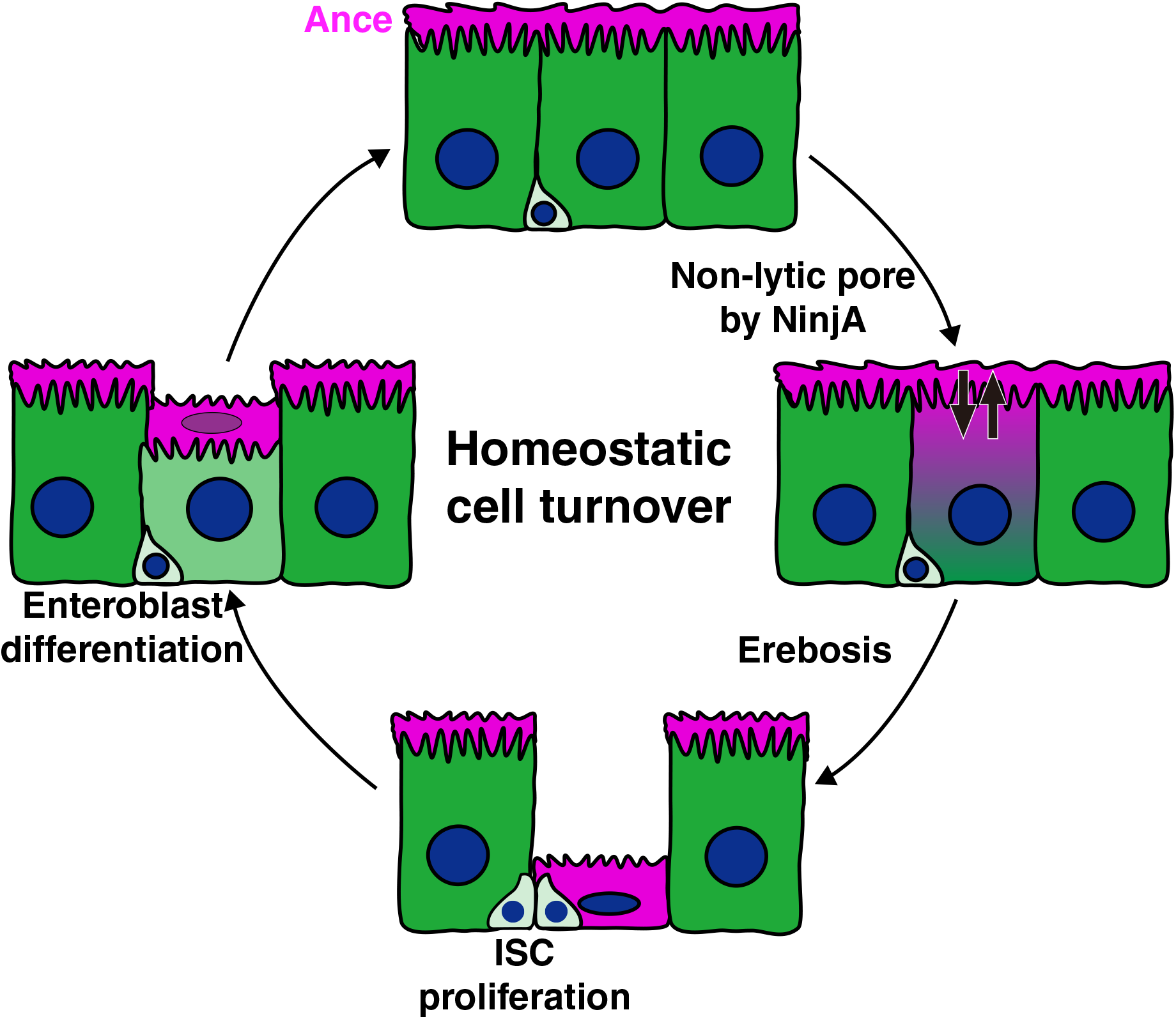
A non-lytic membrane permeabilization by NijA mediates epithelial cell turnover *in vivo*. NijA-mediated pore formation induces erebosis in enterocytes. ISCs located beneath the erebotic cells proliferate and generate new enteroblasts and enterocytes, which push the erebotic cell corpses towards the gut lumen. This cycle supports continuous epithelial turnover in the intestine.

